# Biofilm-forming capacity of *Escherichia coli* isolated from cattle and beef packing plants: relation to virulence attributes, stage of processing, antimicrobial interventions, and heat tolerance

**DOI:** 10.1101/2021.03.24.436903

**Authors:** Kim Stanford, Frances Tran, Peipei Zhang, Xianqin Yang

## Abstract

Despite the importance of biofilm formation in contamination of meat by pathogenic *Escherichia coli* at slaughter plants, drivers for biofilm have been unclear. To identify selection pressures for biofilm, we evaluated 745 ‘Top 7’ from cattle and 700 generic *E. coli* from two beef slaughter plants for motility, expression of curli and cellulose, and biofilm-forming potential. Top 7 were also screened for serogroup, *stx1*, *stx2*, *eae* and *rpoS.* Generic *E. coli* were compared by source (hide of carcass, hide-off carcass, processing equipment) before and after implementation of antimicrobial hurdles. The proportion of *E. coli* capable of forming biofilms was lowest (7.1%; *P* < 0.05) for cattle isolates and highest (87.3%; *P* < 0.05) from equipment. Only one enterohemorrhagic *E. coli* (EHEC) was an extremely-strong biofilm-former, in contrast to 73.4% of *E. coli* from equipment. Isolates from equipment after sanitation had a greater biofilm-forming capacity (*P* < 0.001) than those before sanitation. Most Top 7 were motile and expressed curli, although these traits along with expression of cellulose and presence of *rpoS* were not necessary for biofilm formation. In contrast, isolates capable of forming biofilms on equipment were almost exclusively motile and able to express curli. Results of the present study indicate that cattle would rarely carry EHEC capable of making strong biofilms to slaughter plants. However, if biofilm-forming EHEC contaminated equipment, current antimicrobial hurdles would inadvertently perpetuate the most robust biofilm-forming strains. Accordingly, new and effective anti-biofilm hurdles are required for meat-processing equipment, to reduce future instances of food-borne disease.

**Importance:** As the majority of enterohemorrhagic *E. coli* (EHEC) are not capable of forming biofilms, sources were undetermined of the biofilm-forming EHEC isolated from ‘high-event periods’ in beef slaughter plants. This study demonstrated that sanitation procedures used on beef-processing equipment inadvertently select for survival of the most robust biofilm-forming strains of *E. coli.* Cattle only rarely carry EHEC capable of forming strong biofilms (1/745 isolates evaluated), but sanitation of equipment markedly increased (*P* < 0.001) biofilm-forming capacity of *E. coli*. In contrast, chilling carcasses for 3 days at 0°C reduced (*P* < 0.05) biofilm-forming capacity of *E. coli*. Consequently, an additional anti-biofilm hurdle for meat-processing equipment, perhaps involving cold exposure, is necessary to further reduce the risk of food-borne disease.

## Introduction

High-event periods (HEP) occur sporadically in beef-processing plants when a higher-than expected proportion of trim samples are positive for enterohemorrhagic *E. coli* (EHEC; 1). Isolates of O157:H7 from HEP have a strong ability to form mature biofilms (2), in contrast to the estimated 95% of O157:H7 which lack individual biofilm-forming capacity (3), although EHEC may also become integrated in mixed species biofilms (4). Among non-O157 *E. coli*, biofilm formation is thought to be more common than in O157:H7 (5), but with the exception of O111 and O145 serogroups, carriage of Shiga toxins is also less frequent among non-O157 *E. coli* (6). Impaired biofilm formation in all *E. coli* has previously been linked to three factors: 1) a *stx1* prophage insertion in *mlrA* preventing expression of curli fimbriae; 2) a mutation in *rpoS* reducing expression of both cellulose and curli; or 3) a lack of motility negatively impacting both expression of curli and initial reversible biofilm attachment (7). Biofilm formation among *E. coli* has been thought to be extremely variable among strains (8), but excluding O157, relatively few strains have been evaluated per serogroup (5, 8, 9).

Although isolates from HEP have been characterized, the source of the biofilm-forming *E. coli* causing HEP is not yet determined. One theory is that pathogenic *E. coli* on the hides of cattle contaminate meat products after high bacterial loads overwhelm antimicrobial interventions at one or multiple stages within the processing facility (10). However, as Arthur et al. (1) found little genetic diversity in *E. coli* isolated from HEPs, they proposed that failures in sanitation might occur, transferring a single prevalent strain of EHEC to multiple sites within the slaughter plant. A third possibility, also proposed by Arthur et al. (1), would be that antimicrobial interventions within the slaughter plant would inadvertently select for robust biofilm-forming strains causing HEP. Besides enhanced formation of biofilm, strains of *E. coli* from HEPs have shown increased tolerance to sanitizers (2), although the relationship is unclear for biofilm-forming ability and attributes such as heat resistance which may also allow *E. coli* to survive microbial interventions. Consequently, the present study was undertaken to determine the relationships among biofilm-forming ability of *E. coli* isolated along the production chain including live cattle, hides of carcasses, hide-off carcasses, meat products and processing equipment. For isolates collected in slaughter plants, impacts of antimicrobial interventions (hide wash, carcass chilling, and equipment sanitation) were determined. Factors possibly influencing biofilm-forming ability including seasonality, expression of curli and cellulose, and motility of isolates were evaluated along with the relationship between biofilm-forming capacity and heat resistance. For cattle isolates, impacts of serogroup on biofilm formation and presence of *stx1*, *stx2*, *eae* and *rpoS* were determined.

## Materials and Methods

### Bacterial isolates, antimicrobial hurdles in slaughter plants and culture conditions

*Escherichia coli* isolates (n=1445), included 745 ‘Top 7’ recovered from live cattle or their environment and 700 generic *E. coli* recovered from two federally-inspected beef slaughter plants (A and B) as described by Zhang et al. (11). At plant A, antimicrobial treatments were minimal for carcasses and included carcass trimming, cold-water washing, and air chilling of dressed carcasses. Processing equipment sanitation occurred at the end of each day and included physical removal of detritus, pre-rinsing with pressurized water at 40-50°C, spraying with a chlorine- based alkaline foaming agent, and sanitization with a quaternary sanitizer (12). In contrast, plant B also employed a hide-on carcass wash with 1.5% sodium hydroxide at 55°C, spraying skinned carcasses with 5% lactic acid, and pasteurization of carcasses at the end of the dressing process with steam at > 90°C (13).

Top 7 isolates were screened for serogroup (O26, O45, O103, O111, O121, O145, O157) and the presence of *stx1*, *stx2* and *eae* using primers and PCR conditions described by Conrad et al. (14). Isolates were classified as EHEC if they were positive for *eae* and *stx1* and/or *stx2*. The presence of RNA polymerase sigma factor S (*rpoS)* was detected by PCR as described by Olsen et al. (15). The presence of the locus of heat resistance (LHR) was determined as described by Mercer et al. (16). For generic *E. coli*, their species was verified by PCR using the primers for *uidA* described by Bej et al. (17). After characterization by PCR, each *E. coli* isolate was streaked on MacConkey agar (BD Bioscience, Canada) and incubated at 35°C for 24 ± 2 h. A single colony was then sub-cultured in 5 mL Luria-Bertani (LB; BD Bioscience) and incubated for 16-18 h at 35°C, with shaking at 80-100 rpm.

### Biofilm forming potential

Overnight culture was diluted by taking 50 μl and adding to 5 mL fresh LB, and 160 μl of diluted inoculum was then added into duplicate wells of a round bottom 96-well microtiter plate (ThermoScientific, Canada). Each microtiter plate also included duplicate blank wells of LB as a negative control, while positive controls contained an isolate of O121:H23 known to be a strong biofilm former (18). Microtiter plates were gently covered with a Nunc-TSP 96-peg lid (Fisher Scientific. Canada) to avoid splashing and cross contamination and incubated at 15°C for 4 d. After incubation, the pegged lid was removed and washed with gentle agitation for 1 min in two new Nunc plates each containing 180 μl of phosphate buffer solution (PBS; Millipore Sigma, Canada). The pegged lid was then transferred to a new plate containing 180 μl of 0.1% crystal violet (Millipore Sigma, Canada), stained for 20 min at room temperature with gentle agitation, followed by twice washing the lid in 180 μl PBS with gentle agitation as before. The washed pegged lid was then placed in a new plate containing 180 μl 80% ethanol for 20 min at room temperature with slight agitation.

Absorbance of the microplate at 570 nm was then measured using a microplate reader (Synergy HT, Bio-Tek Instruments Inc., USA). Optical densities (ODs) were then grouped into classes from 0 (non-biofilm former) to 5 (extremely strong biofilm former; Table 1) based on the mean OD of two independent replicates for each isolate as compared to the cut-off OD (ODc) which was equal to three times the standard deviation of OD of negative control plus average OD of negative control.

**Table 1.**
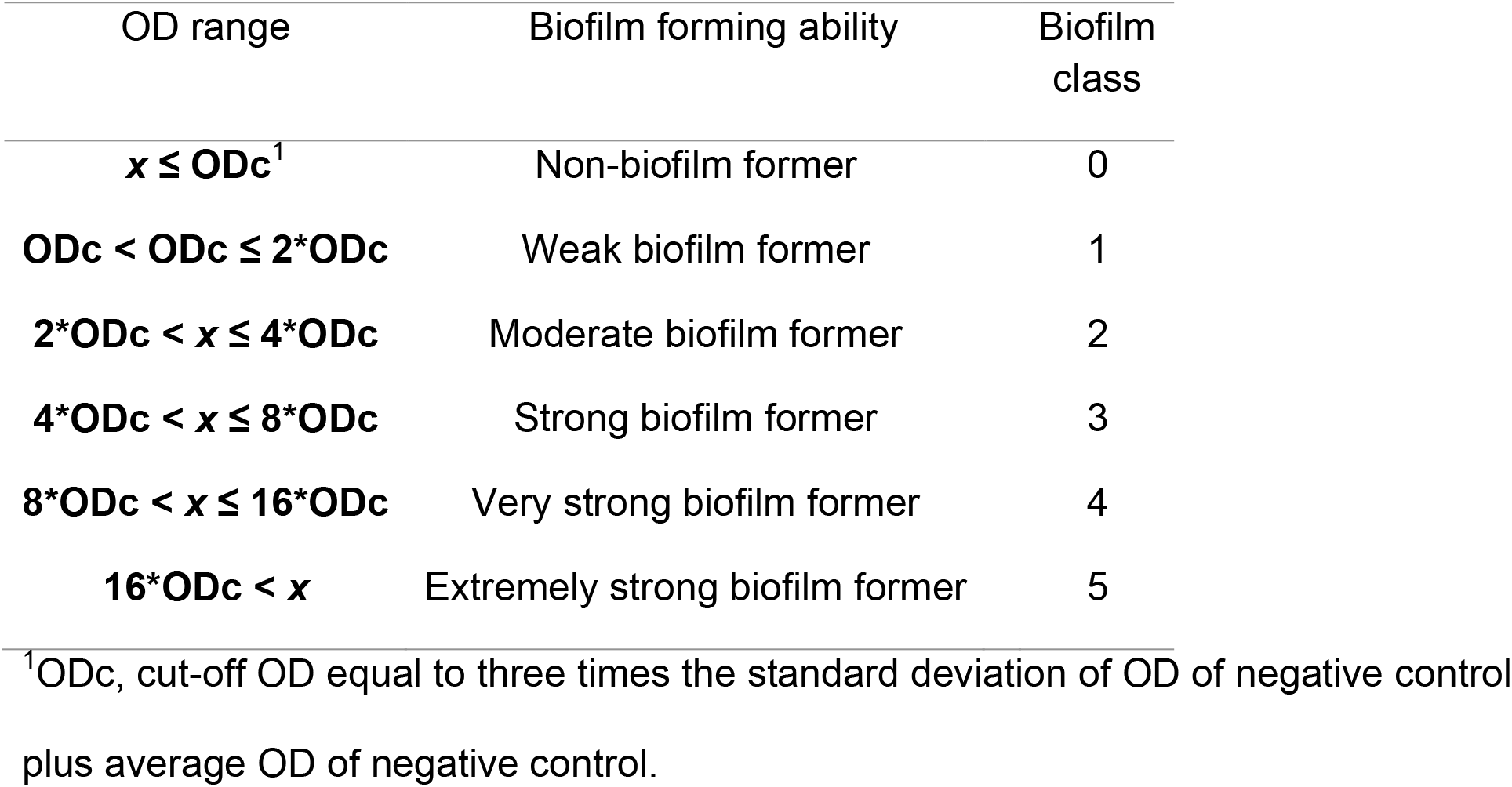
Classification of biofilm-forming ability of isolates based on optical density (OD) at 570 nm of the mean of two independent replicates in duplicate.

| OD range | Biofilm forming ability | Biofilm class |
| --- | --- | --- |
| $x \leq \text{ODc}^1$ | Non-biofilm former | 0 |
| $\text{ODc} < \text{ODc} \leq 2*\text{ODc}$ | Weak biofilm former | 1 |
| $2*\text{ODc} < x \leq 4*\text{ODc}$ | Moderate biofilm former | 2 |
| $4*\text{ODc} < x \leq 8*\text{ODc}$ | Strong biofilm former | 3 |
| $8*\text{ODc} < x \leq 16*\text{ODc}$ | Very strong biofilm former | 4 |
| $16*\text{ODc} < x$ | Extremely strong biofilm former | 5 |

<sup>1</sup>ODc, cut-off OD equal to three times the standard deviation of OD of negative control plus average OD of negative control.

### Motility, expression of curli and cellulose

Motility of isolates was determined using the soft agar method as described by Visvalingam et al. (19). Overnight *E. coli* cultures grown in LB were diluted 100-fold and a 1 μl aliquot was point-inoculated into the center of TYE agar (Tryptone, 10g/; yeast extract, 5g/L; and Bacto agar, 3.5g/L) by stabbing with a micropipette tip halfway through the depth of the agar. Plates were incubated at 15°C for 48 hours. The diameter of each motility halo produced was then measured and an isolate was classified as motile if its halo diameter measured ≥ 4 mm.

The Congo red indicator agar (CRI) method was used to assess the expression of extracellular matrix components (19). Overnight LB cultures were streaked on CRI agar (10 g/L casamino acid, 1 g/L yeast extract, 20 mg/L Congo Red, 10 mg/L Brilliant Blue, and 20 g/L Bacto agar) and incubated at 15°C for 4 days. Color of the resulting colonies determined the phenotype. Colonies were red, brown, pink or white and were respectively considered positive for expression of both cellulose and curli, curli only, cellulose only, or neither (20).

### Phenotypic characterization of heat resistance

Heat inactivation experiments for isolates were as described by Zhang et al. (11). Briefly, 1.5 mL of overnight cultures in LB media in the stationary phase of growth were added to 2mL microcentrifuge tubes (Eppendorf, Canada) and incubated in a 60°C water bath (model FSGPD28, Fisher Scientific^®^, Canada) for up to 30 min. The time required for media to reach 60°C (come up time, T0) was measured by a long needle probe attached to a thermometer (Thermapen^®^ Mk4, Thermoworks, USA) stabbed into the medium through the lids of the tubes. After removal from the water bath, tubes were immediately placed in an ice water bath for rapid cooling. Cultures were then serially diluted in 0.1% (w/v) peptone water and 1ml of appropriate dilutions plated onto Petrifilm Aerobic Count Plates (3M Corp., St. Paul, MN, USA), per manufacturer’s instructions. The plates were incubated at 35°C for 18-24 h and colony forming units (CFU) were enumerated. Log transformed counts were plotted against incubation time and the regression of the plot was used to calculate *D*_60°C_ for each isolate (21).

### Statistical analyses

Statistical analyses were performed using SAS version 9.4 (SAS Institute Inc, Cary, NC). As OD values did not follow normal distributions as determined by Shapiro-Wilk tests, non-parametric analyses (Wilcoxon rank sum test and Kruskal-Wallis test) were used within the NPAR1way procedure to compare the biofilm-forming ability of isolates collected before and after sanitation on OD of *E. coli* sourced from meat processors. For presence/absence values such as formation of biofilm, expression of curli or cellulose, motility and PCR detection of virulence or biofilm-related factors, isolates were compared using generalized linear mixed models (Proc Glimmix) with a binomial distribution. Model adjusted means (back-transformed to original scale) and 95% confidence intervals were determined, with isolate the experimental unit and source of isolate and season fixed effects. Analyses of Top 7 isolates also included serogroup as a fixed effect. To determine relationships between *D*_60°C_ and OD, regression analyses were conducted within Proc Reg for each source of isolates. In all analyses, *P* values < 0.05 were deemed significant.

## Results

### PCR Characterization of Top 7 *E. coli*

Each serogroup differed in the predominant combinations of Shiga toxin genes, *eae* and *rpoS* that were detected (Table 2). For O26, the majority of isolates were *eae* positive and split between those lacking Shiga toxin genes and *rpoS* or EHEC positive for *stx1* and *eae*. In contrast, the largest group of O45 isolates lacked *eae, stx1, stx2* and *rpoS.* Only 13.7% of O45 isolates were EHEC in comparison to 39.2% of O26. Isolates of O103 were evenly split between *eae* positive and negative, with 35.7% EHEC. However, in contrast to O26 and O45, 9.5% of O103 isolates were positive for *rpoS* as compared to only 1% of O45 and 2.9% of O26. Little diversity was present in O111, with 83% EHEC carrying *stx1* and none positive for *rpoS*, although isolate numbers for this serogroup were limited (n=12). Isolates of O121, O145 and O157 were distinguished from other serogroups by more carriage of *stx2*, with 40.2% of O121 isolates EHEC and 16.7% positive for *rpoS.* All isolates of O145 were *eae* and *rpoS* positive, with 46.7% EHEC. By far the largest proportion of EHEC was for O157 (91.2%), with the majority of isolates carrying *rpoS*, *stx1* and *stx2*.

**Table 2.** ‘Top seven’ *E. coli* isolates (n=745) by serogroup and presence of *stx1*, *stx2*, *eae* and r*poS*. Predominant combinations for each serogroup are highlighted in yellow.

| Serogroup<br>and *eae* | No *stx1* |  |  |  |  |  | *rpoS* and |  | Total |
| --- | --- | --- | --- | --- | --- | --- | --- | --- | --- |
|  | *stx2* or<br>*rpoS* | *stx1*<br>only | *stx2*<br>only | *stx1*<br>and *stx2* | *rpoS*<br>and *stx1* | *rpoS*<br>and *stx2* | *stx1* and<br>*stx2* | *rpoS*<br>only |  |
| O26 *eae* + | 45 | 36 | 2 | 1 | 1 | 0 | 0 | 1 | 86 |
| O26 *eae* - | 5 | 5 | 5 | 0 | 1 | 0 | 0 | 0 | 16 |
| O45 *eae* + | 19 | 13 | 0 | 1 | 0 | 0 | 0 | 0 | 33 |
| O45 *eae* - | 43 | 20 | 2 | 3 | 0 | 0 | 0 | 1 | 69 |
| O103 *eae* + | 25 | 33 | 6 | 7 | 2 | 4 | 4 | 6 | 87 |
| O103 *eae* - | 39 | 13 | 0 | 3 | 3 | 0 | 0 | 12 | 70 |
| O111 *eae* + | 0 | 10 | 0 | 0 | 0 | 0 | 0 | 0 | 10 |
| O111 *eae* - | 0 | 2 | 0 | 0 | 0 | 0 | 0 | 0 | 2 |
| O121 *eae* + | 7 | 13 | 15 | 6 | 7 | 0 | 0 | 1 | 49 |
| O121 *eae* - | 23 | 7 | 0 | 14 | 2 | 0 | 0 | 7 | 53 |
| O145 *eae* + | 2 | 0 | 0 | 0 | 13 | 0 | 3 | 17 | 35 |
| O145 *eae* - | 0 | 0 | 0 | 0 | 0 | 0 | 0 | 0 | 0 |
| O157 *eae* + | 1 | 9 | 0 | 6 | 29 | 8 | 162 | 1 | 216 |
| O157 *eae* - | 7 | 8 | 0 | 0 | 2 | 0 | 0 | 2 | 19 |
| **Total** | 216 | 169 | 30 | 41 | 60 | 12 | 169 | 48 | 745 |

### Biofilm-forming potential, motility, expression of curli and cellulose in Top 7

Few Top 7 were able to form biofilm, with 92.9% classified as non-biofilm formers (Table 3). The proportion of EHEC capable of forming biofilms was even smaller, at 2.1%. Only two EHEC capable of forming biofilms also possessed the LHR and were classified as moderate and strong biofilm formers, respectively. As biofilm forming capacity increased, the numbers of Top 7 having these phenotypes decreased, with only two isolates extremely-strong biofilm-formers and one of those EHEC. However, none of the EHEC which were biofilm formers were motile and expressed both curli and cellulose. Overall, motility and expression of curli and cellulose was uncommon in Top 7 with only 25.5% of isolates having all three traits.

**Table 3.**
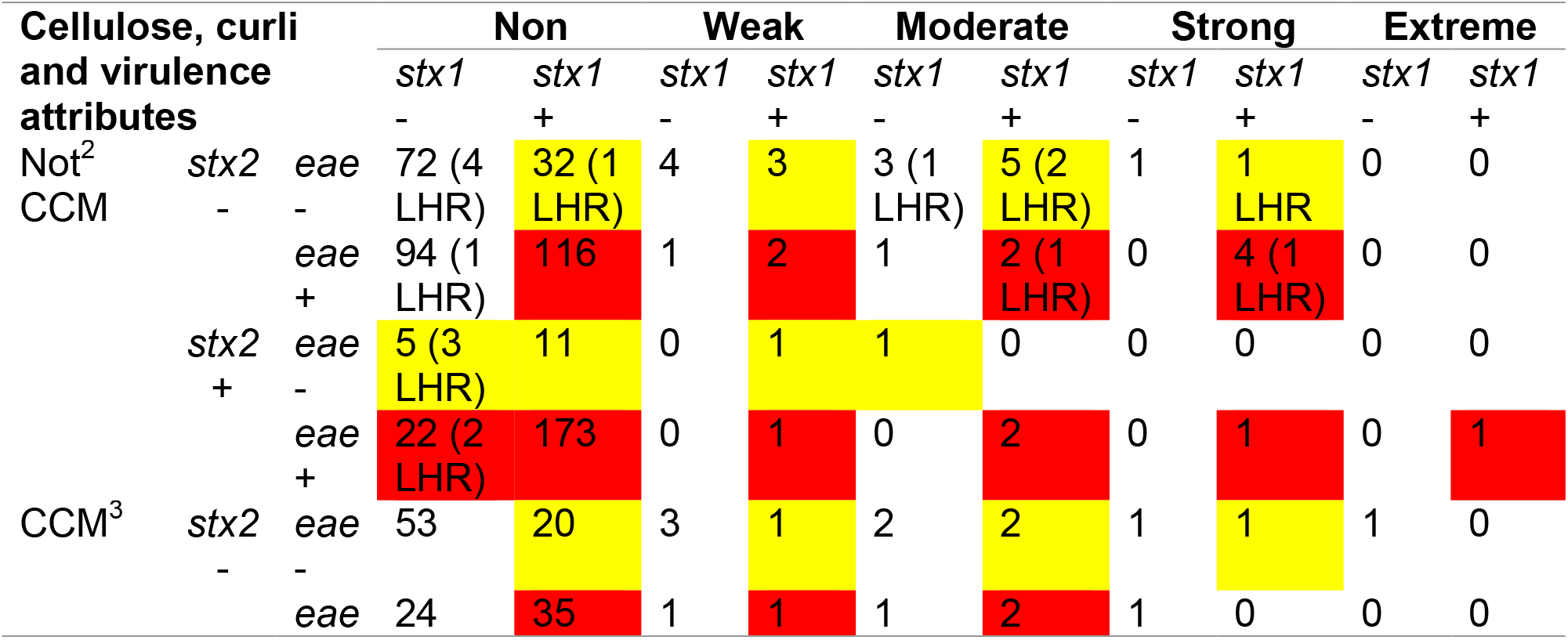
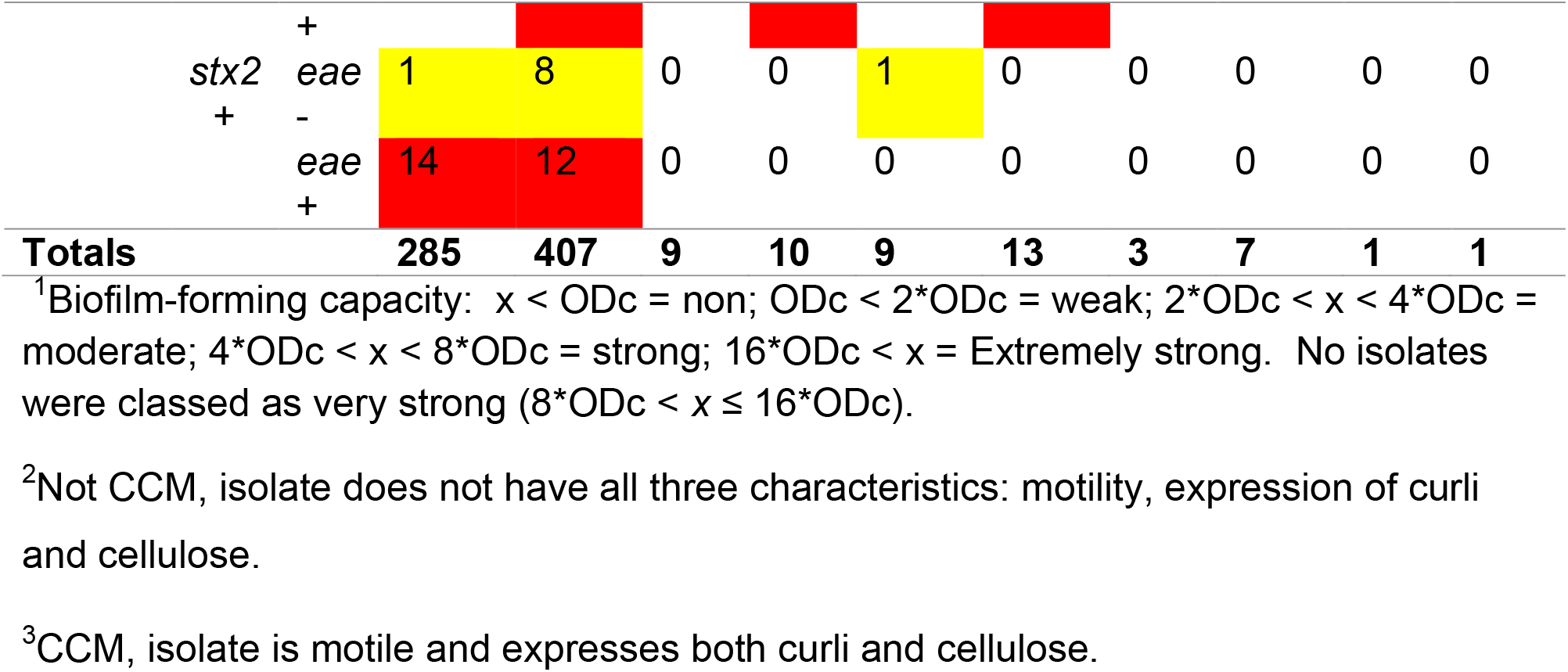
Biofilm forming capacity, curli and cellulose expression, motility, presence of virulence genes and the locus of heat resistance (LHR) in isolates of Top seven *E. coli* (n=745) collected from cattle. Shiga toxin-producing *E. coli* are highlighted in yellow. Enterohemorrhagic *E. coli* are highlighted in red.

Although serogroup did not influence the proportion of Top 7 producing biofilms (Table 4), serogroup did influence motility which was highest (*P* < 0.05) in O157 and lowest in O26, O111 and O145. Similarly, a higher proportion (*P* < 0.05) of O157 isolates expressed curli compared to all other serogroups with the exception of O26. Comparing isolates which both expressed curli and were positive for *stx1*, O157 again had the highest proportion (*P* < 0.05) compared to all serogroups with the exception of O111. However, evaluating the factors that influenced biofilm formation within a serogroup, presence of *eae* and *stx1* negatively influenced (*P* < 0.05) biofilm formation in O157 (Figure 1), meaning that the small proportion of O157 which were not EHEC were generally the strongest biofilm formers. For O157, expression of curli increased (*P* < 0.05) biofilm formation, while expression of cellulose, motility and presence of *rpoS* reduced (*P* < 0.05) biofilm formation. For O26, expression of cellulose increased (*P* < 0.05) biofilm formation, while expression of curli negatively impacted biofilm (*P* < 0.05). For other serogroups, no factors evaluated affected biofilm formation.

**Table 4.**
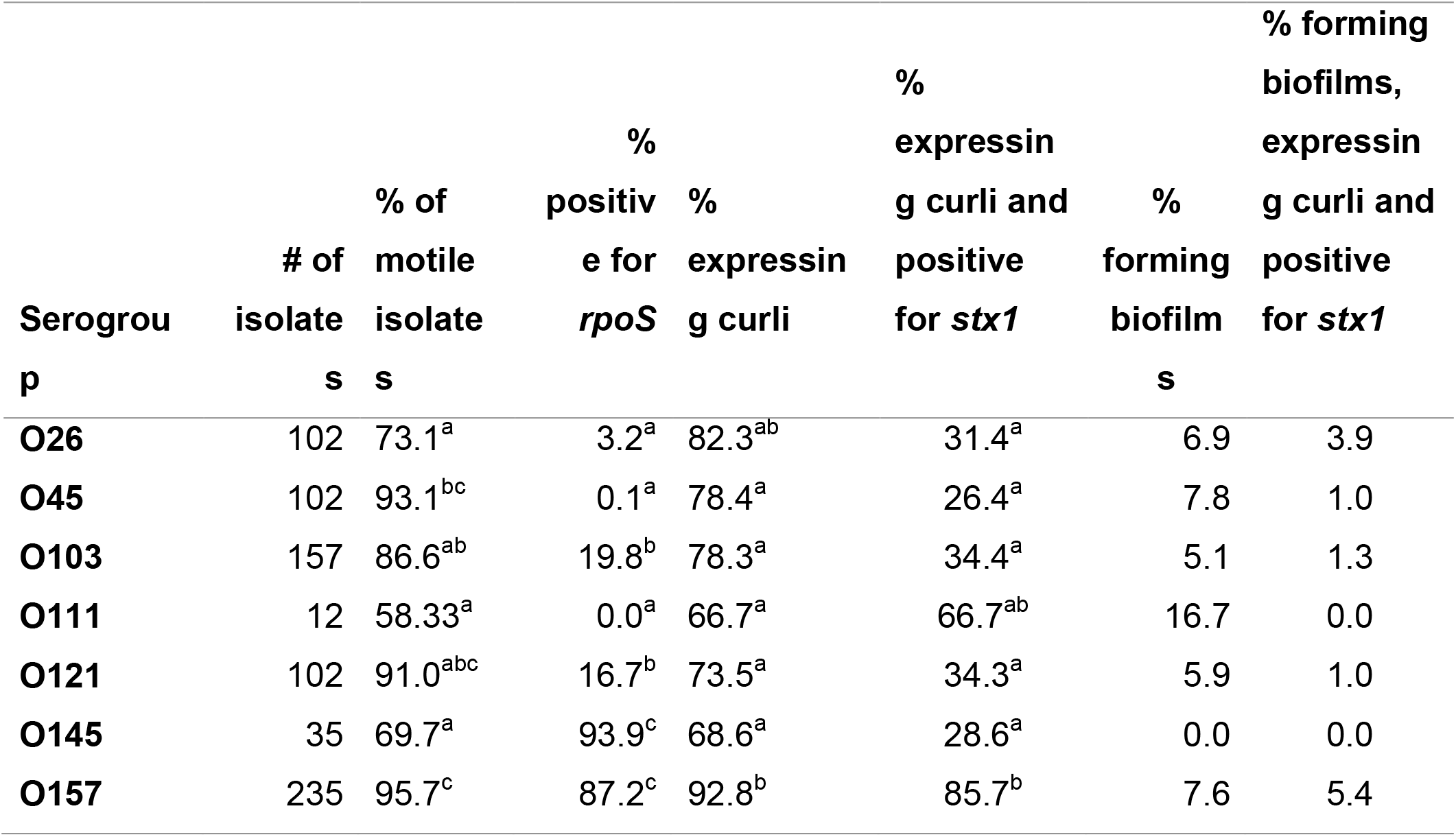
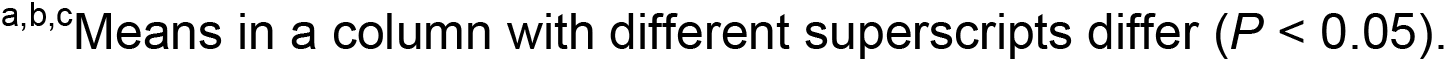
Relationship between expression of curli, presence of *stx1* and production of biofilm by serogroup of Top 7

| Serogroup | # of isolates | % of motile isolates | % positive for *rpoS* | % expressing curli | % expressing curli and positive for *stx1* | % forming biofilm | % forming biofilms, expressing curli and positive for *stx1* |
| --- | --- | --- | --- | --- | --- | --- | --- |
| **O26** | 102 | 73.1<sup>a</sup> | 3.2<sup>a</sup> | 82.3<sup>ab</sup> | 31.4<sup>a</sup> | 6.9 | 3.9 |
| **O45** | 102 | 93.1<sup>bc</sup> | 0.1<sup>a</sup> | 78.4<sup>a</sup> | 26.4<sup>a</sup> | 7.8 | 1.0 |
| **O103** | 157 | 86.6<sup>ab</sup> | 19.8<sup>b</sup> | 78.3<sup>a</sup> | 34.4<sup>a</sup> | 5.1 | 1.3 |
| **O111** | 12 | 58.33<sup>a</sup> | 0.0<sup>a</sup> | 66.7<sup>a</sup> | 66.7<sup>ab</sup> | 16.7 | 0.0 |
| **O121** | 102 | 91.0<sup>abc</sup> | 16.7<sup>b</sup> | 73.5<sup>a</sup> | 34.3<sup>a</sup> | 5.9 | 1.0 |
| **O145** | 35 | 69.7<sup>a</sup> | 93.9<sup>c</sup> | 68.6<sup>a</sup> | 28.6<sup>a</sup> | 0.0 | 0.0 |
| **O157** | 235 | 95.7<sup>c</sup> | 87.2<sup>c</sup> | 92.8<sup>b</sup> | 85.7<sup>b</sup> | 7.6 | 5.4 |

<sup>a,b,c</sup>Means in a column with different superscripts differ ( $P < 0.05$ ).

**Figure 1.**
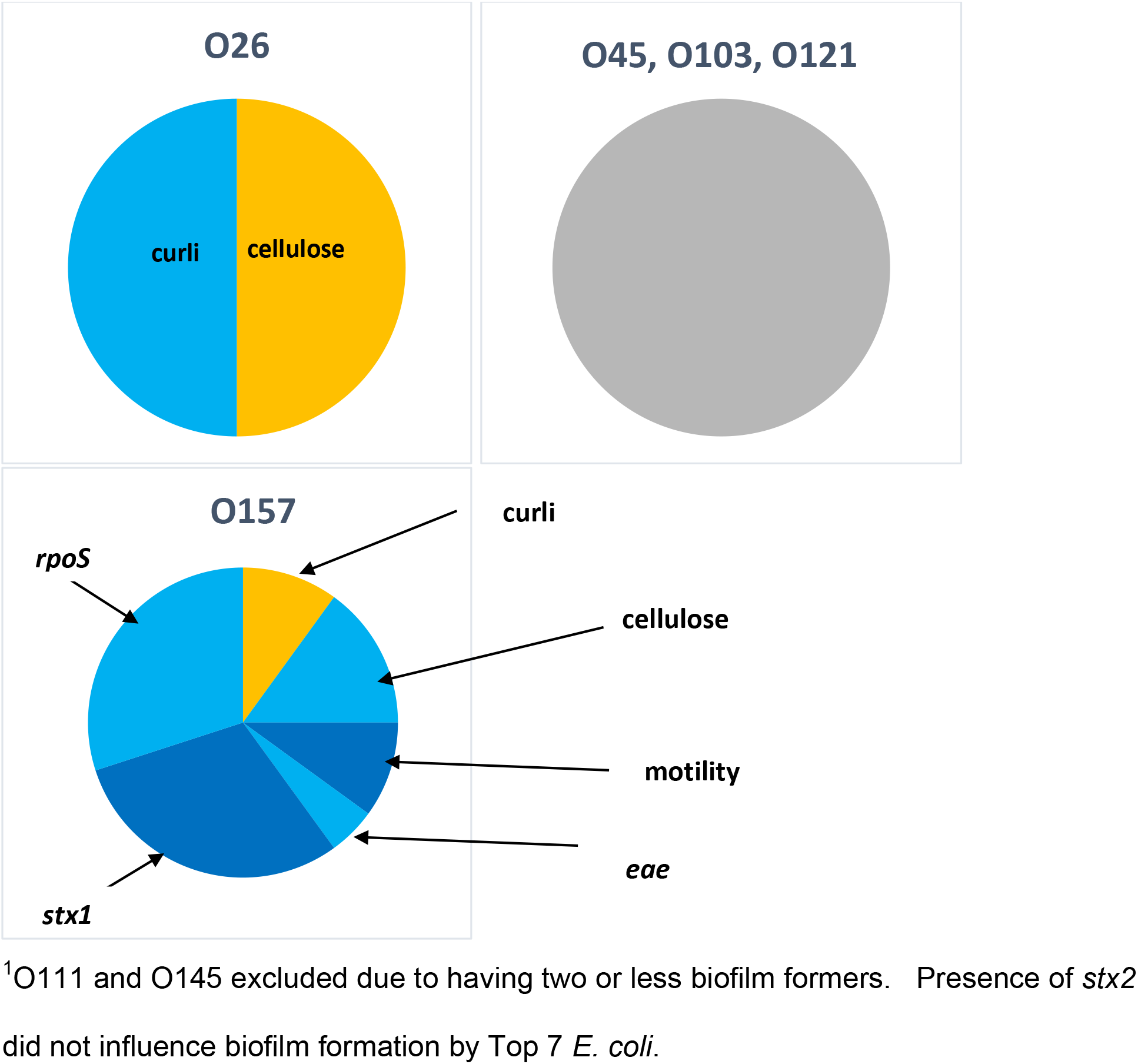
Predictors of biofilm formation by serogroup^1^ of Top 7 *E. coli* including expression of curli and cellulose, motility; presence of *eae*, *rpoS* and *stx1*. Presence of yellow-shaded attributes significantly increased the likelihood of biofilm formation. Presence of light blue or dark blue-shaded attributes significantly reduced the likelihood of biofilm formation. Presence of grey shading indicates no significant predictors of biofilm formation. Proportion of pie is equivalent to relative significance.

### Biofilm-forming potential, motility, expression of curli and cellulose in generic *E. coli*

In contrast to Top 7, 42.9% of generic *E. coli* were capable of forming biofilms (Table 5). From processing equipment, 70% of isolates were strong biofilm formers which were motile and expressed both curli and cellulose. At the opposite extreme, 73% of isolates from beef products were non-biofilm formers, with 40% of beef isolates motile and expressing both curli and cellulose. Hide-off carcass isolates were 53.0% biofilm formers, and 49.8% were motile expressing curli and cellulose. Similar to hide-off carcasses, isolates collected from the hide of carcasses were 44.6% biofilm formers, with 42.1% motile and expressing both curli and cellulose. A total of seven isolates collected from hide-off carcasses classified as very-strong or extremely-strong biofilm-formers also possessed the LHR, although no LHR-positive isolates were present in beef products or from processing equipment.

**Table 5.** Biofilm-forming capacity, presence of the locus of heat resistance (LHR), motility, curli and cellulose expression by stage of processing (hide of carcass, hide-off carcass, beef product; n=500) or processing equipment (n=200) for generic *E. coli* isolates collected in two federally-inspected beef slaughter plants in Alberta.

| Biofilm-forming<br>Capacity<sup>1</sup> | Hide of carcass |  | Hide-off carcass |  | Beef Products |  | Processing<br>Equipment |  |
| --- | --- | --- | --- | --- | --- | --- | --- | --- |
|  | MCC<sup>2</sup> | Not<br>MCC<sup>3</sup> | MCC | Not MCC | MCC | Not<br>MCC | MCC | Not MCC |
| **Non** | 21 | 46 (1<br>LHR) | 54 | 77 | 34 | 39 | 24 | 5 |
| **Weak** | 1 | 3 | 13 | 10 | 1 | 5 | 4 | 0 |
| **Moderate** | 4 | 5 | 5 | 4 | 1 | 8 | 13 | 1 |
| **Strong** | 8 | 5 | 10 | 22 | 2 | 4 | 4 | 5 |
| **Very**<br>**strong** | 5 | 3 | 25 | 19 (5<br>LHR) | 1 | 3 | 5 | 0 |
| **Extreme** | 12 | 8 | 32 | 8 (2 LHR) | 1 | 1 | 138 | 1 |
| **Totals** | **51** | **70** | **139** | **140** | **40** | **60** | **188** | **12** |

<sup>1</sup>Biofilm-forming capacity: $x < \text{ODc}$ = non; $\text{ODc} < 2*\text{ODc}$ = weak; $2*\text{ODc} < x < 4*\text{ODc}$ = moderate; $4*\text{ODc} < x < 8*\text{ODc}$ = strong; $8*\text{ODc} < x < 16*\text{ODc}$ = very strong; $16*\text{ODc} < x$ = Extremely strong.
<sup>2</sup>Isolate is motile and expresses both curli and cellulose.
<sup>3</sup>Isolate does not have all three characteristics: motility, expression of curli and cellulose.

### Effects of source of isolates and seasonality on biofilm formation

Comparing all sources, the proportion of isolates capable of forming biofilm was highest (*P* < 0.05) in those collected from processing equipment, lowest (*P* < 0.05) in those from live cattle, and intermediate in isolates from hides of carcass, hide-off carcass and beef products (Figure 2). Significant interactions between source of isolates and two biofilm-related phenotypes (expression of curli and motility) were present (Figure 3). While expression of curli had no influence on the proportion of live cattle or beef product isolates making biofilms, this phenotype increased (*P* < 0.05) the proportions of isolates from processing equipment and from hides of carcasses which formed biofilms. Unexpectedly, in hide-off carcasses, expression of curli reduced (*P* < 0.05) the proportion of isolates forming biofilms. For both hide-off carcasses and processing equipment, motility increased (*P* < 0.05) the proportion of isolates forming biofilms, but did not influence biofilm formation in isolates from live cattle, hides of carcasses or beef products. For isolates from live cattle, hides of carcasses and hide-off carcasses where season of isolate collection was known, biofilm forming potential was increased (*P* < 0.05) in the winter as compared to the fall, with summer months intermediate (Figure 4).

**Figure 2.**
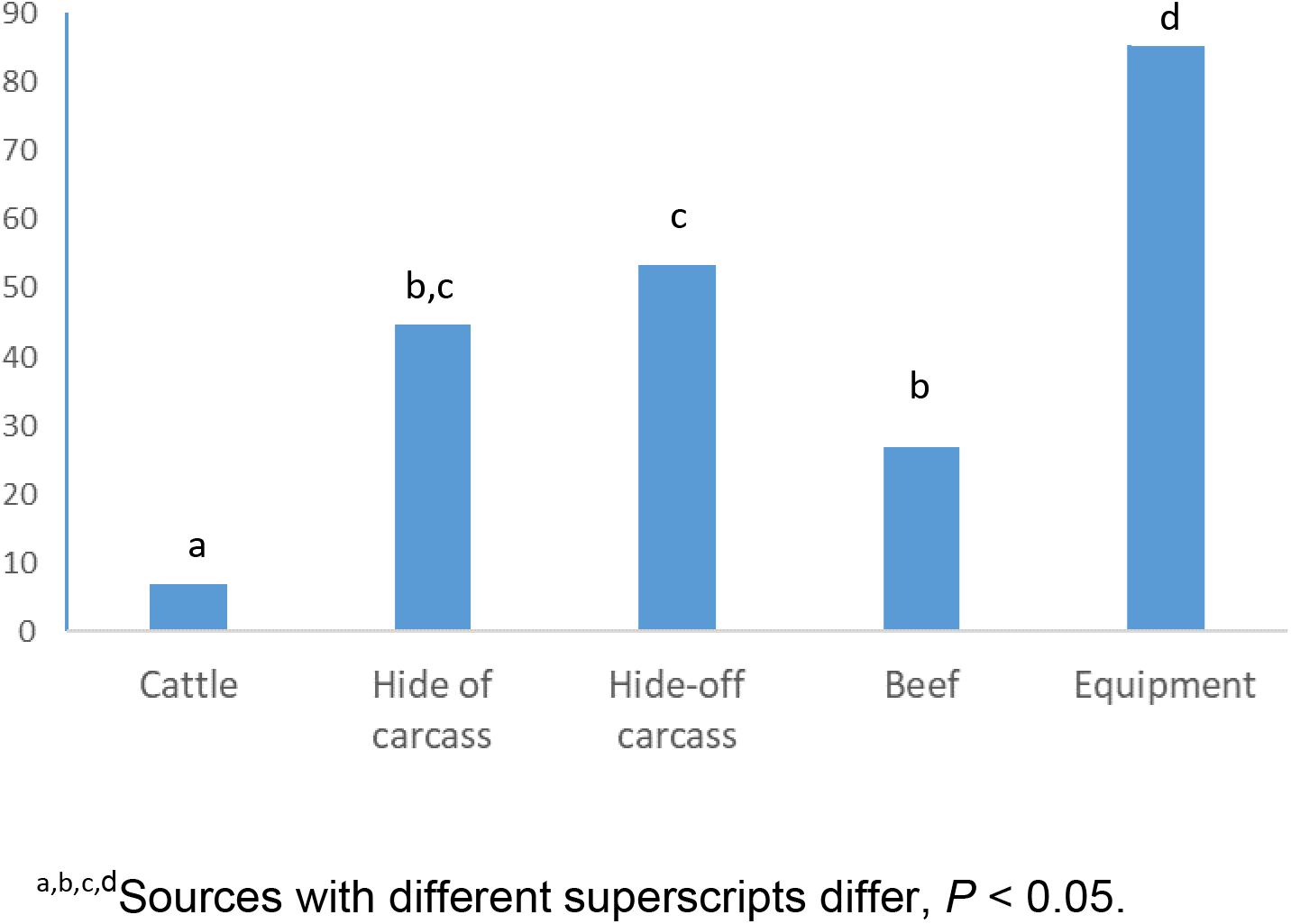
Effect of source on proportions of *E. coli* capable of forming biofilms as determined by optical density at 570 nm.

**Figure 3.**
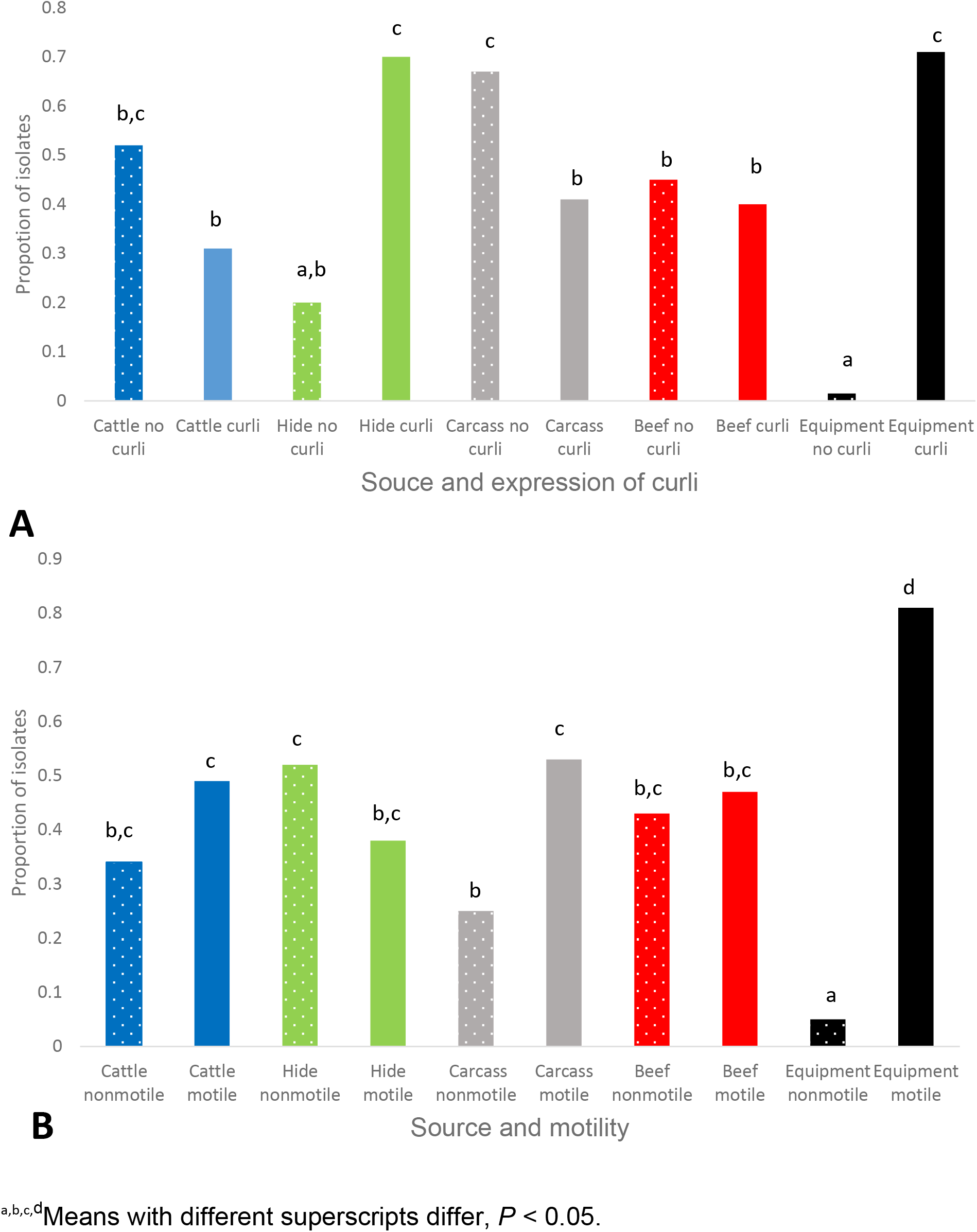
Interactions between source and phenotypic attributes for proportions of isolates of *E. coli* forming biofilms as determined by optical density (OD) at 570 nm: expression of curli (A), motility (B).

**Figure 4.**
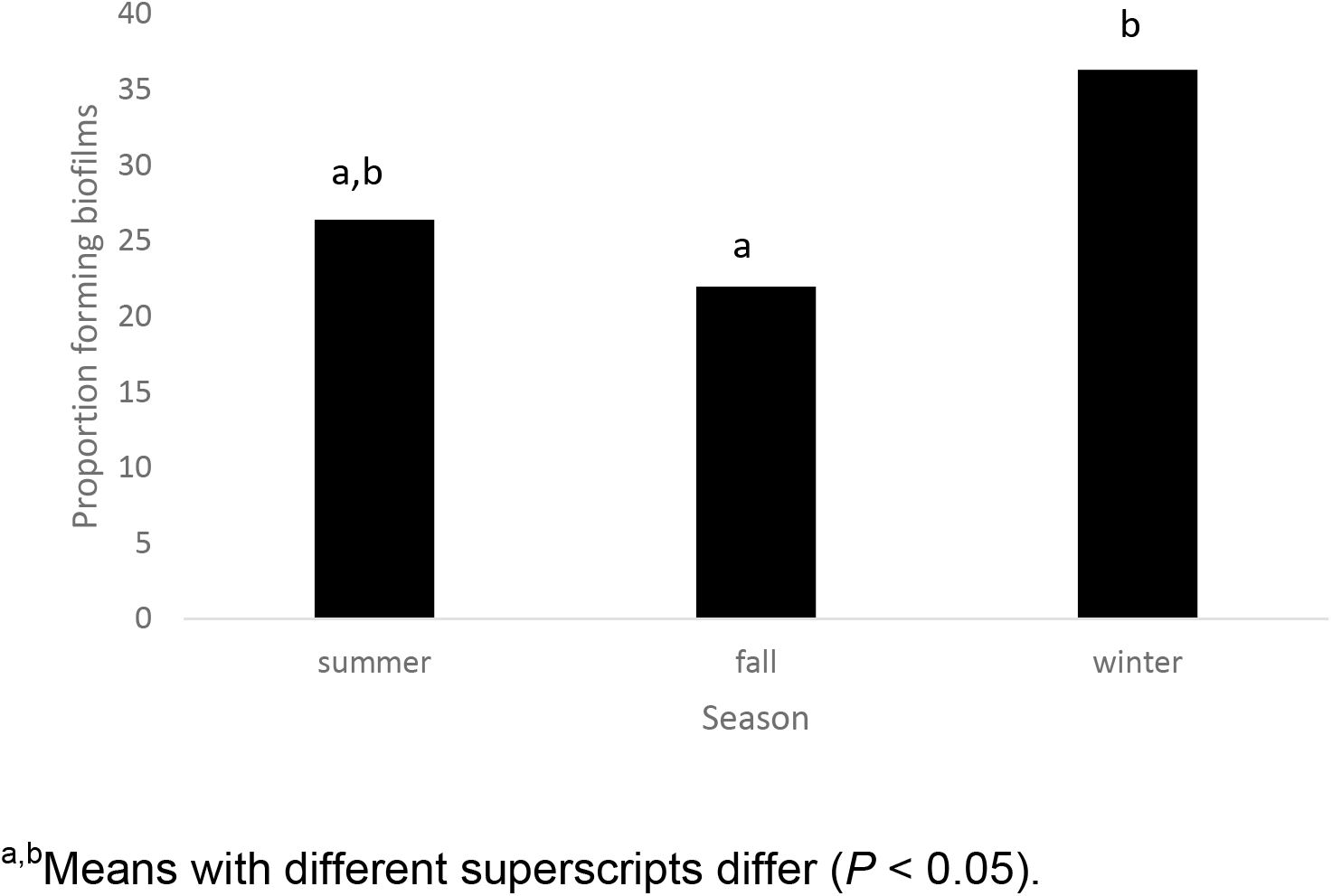
Proportion of isolates forming biofilms by season. Includes *E. coli* isolated from cattle, hides of carcasses and hide-off carcasses (n=658).

### Impacts of antimicrobial interventions at slaughter plants on biofilm

Of the three antimicrobial interventions evaluated, hide washing was the only one which did not significantly impact subsequent biofilm-forming capacity of isolates (Figure 5). Chilling reduced (*P* = 0.018) biofilm formation by isolates collected after carcasses were held at 0°C for 3 d. In contrast, sanitation of equipment exerted a strong selection pressure for biofilm, with biofilm-forming capacity of isolates remaining after sanitation markedly increased (*P* < 0.001).

**Figure 5.**
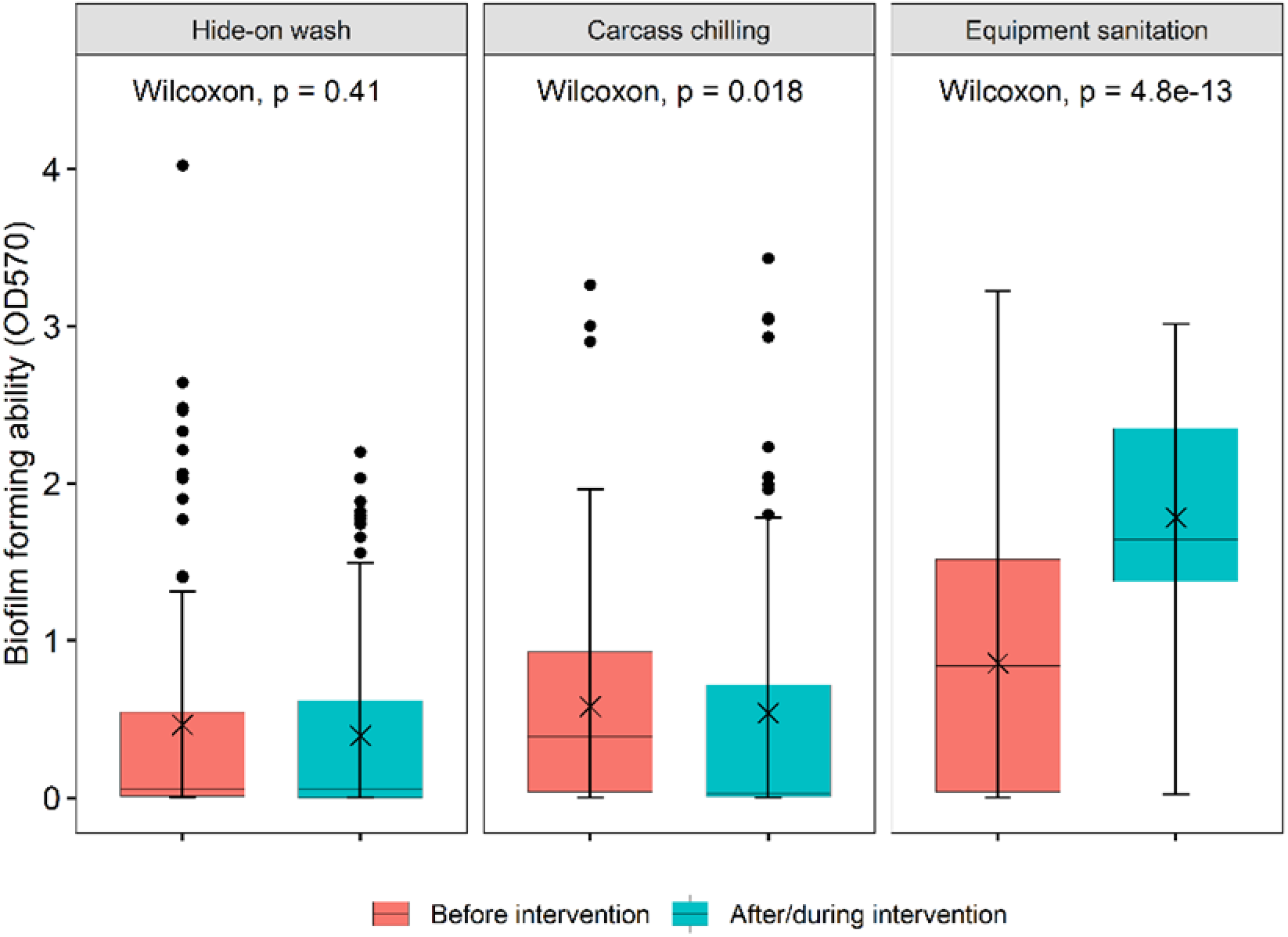
Effect of antimicrobial interventions at beef processing plants on biofilm-forming ability of *E. coli* isolates as determined by optical density (OD) at 570 nm.

### Relationship between biofilm formation and heat resistance

Although biofilm-formation as measured by OD and heat resistance of isolates as determined by D_60oC_ were not significantly related in regression analyses (Figure 6), heightened heat resistance and biofilm-forming ability were not mutually-exclusive. With the exception of live cattle, all sources had one or more isolates which were relatively heat-resistant biofilm-formers. Presence of the LHR did not influence heat resistance of these biofilm-forming isolates as no isolates from processing equipment were positive for the LHR, yet four of these were both heat resistant (D_60oC_ > 2 min) and extremely strong biofilm formers. Similarly, although two strong biofilm formers isolated from live cattle were positive for the LHR, D_60oC_ for these isolates were < 1.2 minutes, similar to the majority of cattle isolates.

**Figure 6.**
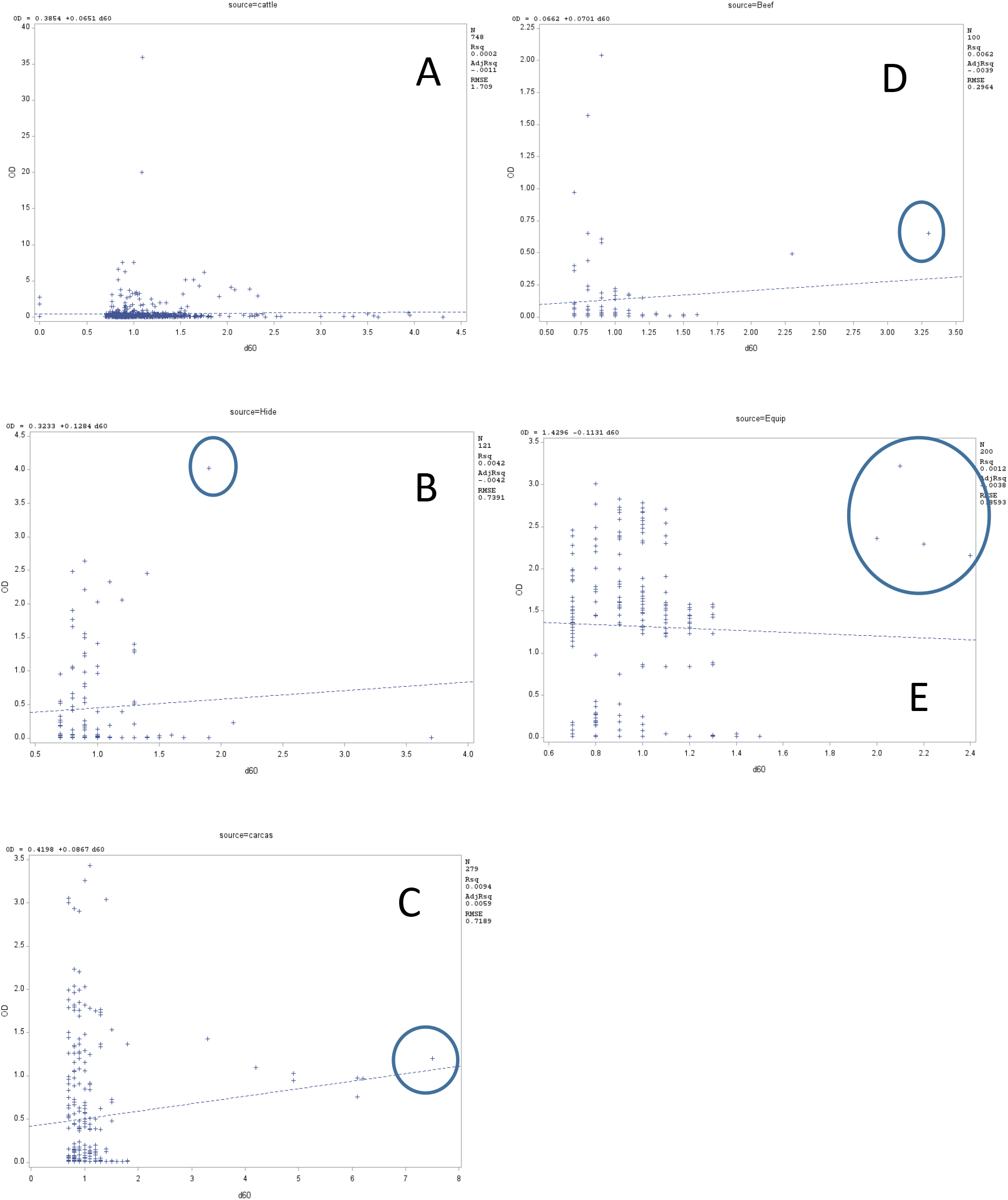
Relationship between biofilm optical density and heat tolerance (D_60oc_) of *E. coli* by source of isolates: live cattle (A), hide of carcass (B), hide-off carcass (C), beef products (D), processing equipment (E). Isolates having relatively heightened heat resistance and biofilm forming capacity are circled.

## Discussion

### Biofilm formation by Top 7

Although it has been reported that 95% of O157 strains are unable to express curli (22), this was not the case in the present study and the majority of Top 7 shared this capability. Prophages carrying *stx1* have been found to insert within and inactivate *mlrA*, which is a positive regulator for CsgD and necessary for the transcription of curli operons (3). However, in reverse of expectations, expression of curli and carriage of *stx1* were higher in O157 compared to most serogroups evaluated. Some curli-positive *stx1*-positive O157 strains have a mutation in the *csgD* promoter which leads to over-expression of curli, although other curli-positive O157 strains have been *stx1*-negative after loss of the *mlrA*-embedded prophage (23). Whole genome sequence analyses of curli-expressing *stx1-*positive O157 are in progress and may better explain our results. Potentially, prophage carrying *stx1* may integrate at sites other than within *mlrA*, or mutations in the *csgD* promotor may be more common than previously thought, as curli-expressing *stx1*-positive isolates were more than 25% of those evaluated for all serogroups of Top 7.

Even though curli was expressed by the majority of Top 7, few were capable of forming biofilms. Strains of EHEC that do not produce curli or cellulose and are non-motile are thought to have less biofilm-forming capacity than strains with a combination of these traits (9). Flagella have been shown to be important in the initial attachment of *E. coli* to a surface (24), but in the present study expression of cellulose was the only trait positively influencing biofilm formation by O26, while expression of curli was the only trait positively influencing biofilm formation by O157 (Figure 1). Motility and expression of cellulose were actually negatively associated with biofilm formation by O157, but these and others such as carriage of *rpoS*, and *eae* may have been negatively associated only due their predominance in EHEC. Even though non-EHEC O157 strains were uncommon, they constituted 29% of O157 biofilm-formers. Although not all isolates were serotyped, some of the strongest biofilm-forming O157 were O157:H12, which has been used as a non-pathogenic surrogate for *E. coli* O157:H7 (25). Even though EHEC forming biofilms were rare in the present study, and similar to previously reported O157:H7 biofilm-forming prevalence (3), they were not unknown. One EHEC was an extremely-strong biofilm former (Table 3) as has been reported for EHEC responsible for HEP at abattoirs (2).

All Top 7 were evaluated for presence of r*poS* due its role in improved tolerance to stress by O157:H7 as well as its influence on biofilm formation as the master regulator of the ‘curli/cellulose control cascade’ (24). Results of the present study, however did not demonstrate any positive role of *rpoS* in biofilm formation, possibly due to incubation conditions used. Although biofilms were evaluated in stationary phase, *rpoS* is expressed during periods of stress and starvation (26), which may not have occurred in incubations in a nutrient-dense medium such as LB. No biofilm was formed by isolates of O145 although it along with O157 had the highest (*P* < 0.05) proportions of *rpoS* detected (Table 4). Potentially, *rpoS* may have been present, but not expressed and genomic evaluations currently in progress will determine if mutations in *rpoS* may be leading to a lack of function. Both motility and expression of cellulose were lower (*P* < 0.05) in isolates of O145 as compared to O157, which perhaps contributed to less biofilm formation by O145, although motility was also not positively linked to biofilm formation by Top 7 in the present study.

Picozzi et al. (8) concluded that biofilm formation among EHEC was extremely variable and could not be predicted by serogroup or by presence of virulence genes, although they evaluated only 45 isolates across six serogroups. Our results demonstrate that other than by directly evaluating biofilm formation, it was not possible to predict which Top 7 isolates would be biofilm formers based on motility, expression of curli and cellulose, and presence of *rpoS*, in agreement with a recent study by Ma et al. (27) which concluded that genetic or phenotypic characterizations of curli, cellulose, and motility did not correspond with biofilm-forming potential of *E. coli*. These authors also evaluated an extensive list of 21 biofilm-related genes, the presence or absence of which were also not able to predict biofilm formation. Based on our results none of the three factors identified by Chen et al. (7) as reasons for impaired biofilm formation could be confirmed in the Top 7 evaluated, as motility and expression of curli and cellulose were common compared to their capacity to form biofilm. Methods to quantify expression of curli and cellulose may provide additional insight into biofilm formation by EHEC.

### Effects of source of isolates and antimicrobial interventions on biofilm formation

To our knowledge, this would be the first study to compare biofilm-forming capacity of *E. coli* isolated from the complete beef production chain. In contrast to our previous study of the same isolates, which for heat tolerance found no differences among sources of isolates or evidence of antimicrobial interventions exerting a selection pressure for increased heat tolerance (11), both source of isolates and antimicrobial interventions affected biofilm-forming capacity of isolates. Comparing sources, heightened biofilm-forming capacity was found in isolates from processing equipment compared to all others evaluated. As well, sanitation of processing equipment was confirmed to be a strong selective pressure for increased biofilm. Previous work raised this possibility as biofilm-forming ability was found to improve persistence of *E. coli* on meat fabrication equipment (28). In the present study, only strong biofilm formers were likely able to remain on equipment and be recovered after sanitation. As chilling at 0°C reduced biofilm formation on carcasses, perhaps a chilled water wash or other chilling procedure might be a beneficial additional anti-biofilm hurdle after completion of equipment sanitation, although additional studies would be necessary to develop and evaluate efficacy of such a process, and Dourou et al. (29) found that biofilms of O157:H7 could form on beef processing surfaces at 4°C.

Other than being numerous, biofilm-forming isolates from equipment were also notable as they were almost exclusively motile (Figure 3), although motility also increased (*P* < 0.05) the proportion of biofilm formers on carcasses. Motility of isolates has been often linked to the ability of *E. coli* to form biofilms (7, 27, 30) and while not associated in the present study with biofilm formation in isolates from cattle, the hides of carcasses or meat products, motility may be more necessary to form biofilms on processing equipment and hide-off carcasses. Perhaps movement to locations more protected from sanitation procedures affecting carcasses and processing equipment are a requirement for biofilm initiation and overall survival of *E. coli* from these sources. Similarly, Chitlapilly Dass et al. (30) theorized that motile, biofilm-forming EHEC swim upstream after sanitation and accumulate in protected locations such as floor drains of slaughter facilities.

Along with requirements for motility, biofilm-forming isolates from processing equipment also almost exclusively expressed curli, a phenotype shared by the majority of biofilm-forming isolates from the hides of carcasses. Both hides and processing equipment would be routinely subjected to high-pressure water washes in contrast to other sources of *E. coli* evaluated, with the exception of hide-off carcasses. That the proportion of isolates from hide-off carcasses which formed biofilm was reduced in those expressing curli was unexpected, but a number of adhesins contribute to biofilm formation (27) and alternatives to curli may be more important for adherence of *E. coli* to the layer of subcutaneous fat on the surface of the hide-off carcass.

### Seasonality of biofilm formation

Seasonality of *E. coli* biofilm formation in the beef production chain has been little examined, although several studies have evaluated seasonality of multi-species biofilms in rivers or water systems (31,32). Although seasonality results of the present study would be preliminary as data were not available for all four seasons, results were consistent for isolates from cattle, hides of carcasses and hide-off carcasses, the only isolates where season of collection was known. Seasonal differences in isolates would be most expected in those collected from cattle and their environment as many were collected from fecal pats or the hides of cattle exposed to ambient temperatures.

Biofilm forming ability was heightened in the winter when temperatures in Alberta are commonly < 0°C (33). Genetic diversity of *E. coli* collected from cattle in winter months has been shown to be reduced (6), likely as only a smaller, more-related population are able to survive. With surviving populations in the winter likely cold-stressed, and stress linked to expression of *rpoS* and production of curli and cellulose (26), perhaps the increased proportion of biofilm-formers in the winter is related to increased stress to *E. coli*. The close relationship among Top 7 isolates from cattle and generic *E. coli* collected at slaughter plants has been previously demonstrated (11), with increased biofilm-forming capacity of cattle isolates entering the slaughter plant during the winter likely passed to other steps of the slaughter chain. Although seasonality of biofilm-formation on processing equipment is as yet unknown, any increased biofilm-forming capacity in the winter would be balanced by a reduced population of viable *E. coli* (6), potentially reducing the risk of transfer of EHEC to processing equipment.

### Relationship between biofilm-forming ability and heat resistance

Biofilm-forming capacity of *E. coli* is known to be temperature-dependent (8, 29), but it is only recently that the relationship between heat resistance and biofilm formation was investigated by Ma et al. (27). These authors found that the six heat-resistant (LHR- positive) generic *E. coli* that they investigated were all biofilm-formers. While we would agree with these authors that heat-resistant biofilm-forming strains may be a ‘serious food safety and public health risk’, evaluating larger numbers of *E. coli* found no relationship between the LHR and biofilm forming capacity, or between heat resistance as determined by D_60oC_ and OD of biofilms formed. Of 24 isolates evaluated with the LHR, 54% were capable of forming biofilms, with the majority of these generic *E. coli.* Instead, results of the present study would support that even on processing equipment where biofilm-formers were most common, heat-resistant biofilm-forming isolates (n=4, none with the LHR) were rare.

### Most likely mechanisms for HEP

Arthur et al. (1) postulated that failures in hygiene may spread a single dominant strain of EHEC throughout a slaughter plant or that sanitation procedures within the slaughter plant were perhaps inadvertently selecting for strains of EHEC capable of causing HEP. Results of the present study would support roles for both of these mechanisms in establishment of biofilm-forming EHEC capable of causing a HEP. Biofilm-forming EHEC were rare and only 1/745 *E. coli* isolated from cattle was an EHEC and an extremely strong biofilm-former. As a strong-biofilm former, potentially this or other similar EHEC would have the highest likelihood of surviving sanitation procedures and contaminating processing equipment, perhaps being transported through wash water pipes as proposed by Chitapilly Dass et al. (30) to multiple areas of the slaughter plant. If EHEC biofilm-formers were to contaminate processing equipment, results of the present study conclusively demonstrate that current equipment sanitation procedures would exert a strong selection pressure for the most robust biofilm formers. Accordingly, a new and effective anti-biofilm hurdle is required for processing equipment if we are ever to reduce HEP and increase the safety of beef products.

## Acknowledgements

Many thanks to the Beef Cattle Research Council for project funding (FOS.01.17) and to Susanne Trapp, Allison McNaughton, and Rebecca Lohmann for sterling technical assistance.

